# Global optimization approach for circular and chloroplast genome assembly

**DOI:** 10.1101/231324

**Authors:** Sebastien François, Rumen Andonov, Dominique Lavenier, Hristo Djidjev

## Abstract

We describe a global optimization approach for genome assembly where the steps of scaffolding, gap-filling, and scaffold extension are simultaneously solved in the framework of a common objective function. The approach is based on integer programming model for solving genome scaffolding as a problem of finding a long simple path in a specific graph that satisfies additional constraints encoding the insert-size information. The optimal solution of this problem allows one to obtain new kind of contigs that we call distance-based contig. We test the algorithm on a benchmark of chloroplasts and compare the quality of the results with recent scaffolders.

## 1 Introduction

Modern Next-Generation Sequencing (NGS) techniques output billions of short DNA sequences, called *reads*, and the typical way to process this information is by using *de novo* assembly. However, assembling these fragmented raw data into complete genomes remains a challenging computational task. This is a very complex procedure, usually involving three main steps: (1) generation of *contigs*, which are contiguous genomic fragments issued from the overlapping of the reads; (2) constructing *scaffolds*—sequences of oriented contigs along the genome interspaced with gaps; (3) *finishing*, which aims to complete the assembly by inserting DNA text in the gaps between the ordered contigs.

The first step generates a list of contigs (also called *unitigs*) that usually represent the “easily assembled regions” of the genome. Building contigs is currently supported by methods using a specific data structure called *de-Bruijn* graph [13], where genomes are sought as maximal unambiguous paths. Despite the progress done by the community in the domain, complex regions of the genome (e.g., regions with many repeats) generally fail to be assembled by these techniques. If the genome contains repeats longer than the size of the reads, the entire genome cannot be built in a unique way.

Whereas the main challenge of the first step relies on handling huge volume of data, the scaffolding step manipulates data of moderate size. However, the problem remains largely open because of its NP-hard complexity [9]). The goal here is to provide a reliable order and orientation of the contigs in order to link them together into *scaffolds*. Contigs can be linked together using *paired-end* or *mate-pair* reads [16, 11]. This complementary data is due to the ability of the sequencing technology to provide couples of reads that are separated by a known distance (called *insert size*). They bring a long distance information that is not used in the first assembly stage, but is essential for the second.

The scaffolding phase usually produces multiple scaffolds. Moreover, these scaffolds may contain regions that have not been completely predicted. Hence, two additional steps, *gap-filling* and *scaffold extension* (elongating and concatenating the contigs after the scaffolding step) are typically needed to complete the genome.

The strategy proposed here differs significantly from the approaches described in the literature. While the latter apply various heuristics for tackling the different assembly stages one after another separately, our methodology consists of developing a global optimization approach where the scaffolding, gap-filling, and scaffold extension steps are simultaneously solved in the framework of a common objective function. Our approach is based on integer programming models for solving the genome scaffolding as a problem of finding a long simple path in a specific graph that satisfies additional constraints encoding the insert-size information [4].

We are not aware of previous approaches on scaffolding based on longest path problem reduction. Most previous work on scaffolding is heuristics based, e.g., SSPACE [2], GRASS [5], BESST [15] and SPAdes [1]. Such tools may find in some cases good solutions, but their accuracies cannot be guaranteed or predicted. Exact algorithms for the scaffolding problem are presented in [17], but the focus of that work is on finding structural properties of the contig graph that will make the optimization problem of polynomial complexity. In [12], integer linear programming is used to model the scaffolding problem, with an objective to maximize the number of links that are satisfied. In order to avoid sub-cycles in the solution, the authors use an incremental process, where cycles that may have been produced by the solver are forbidden in the next iteration. Integrating the distances between contigs and accounting for possible multiplicities of the contigs (repeats) is indicated as future improvement in [12], while it has been realized in our approach.

This paper focuses on circular genomes and, in particular, on chloroplasts. The reasons for this choice are as follows. Chloroplasts possess circular and relatively small genomes. The particularity of these genomes is the presence of numerous repetitions, while these are the main chalenges for the modern genome assembly techniques. On the other hand, the size of the chloroplast genome permits assembling them rapidly (each one of the instances from the considered benchmark except one, EuglenaGracilis genome, has been solved for less that 1 sec.) and so we were able to refine our strategy and to focus entirely on the quality of the obtained results.

The contributions of this study are as follows:

- We adapt and further develop the general case approach proposed in [4] to the case of circular genomes. Using the specificities of this particular case we succeed to simplify significantly the sophisticated mixed integer linear program (MILP) described in [4].
- We propose an exact approach for scaffolding in the case of circular genomes as a problem of finding longest paths in specific unitig graphs with additional set of constraint distances between couples of vertices along these paths.
- We deeply analyze the reasons for the existence of a huge number of multiple equivalent optimal solutions. These solutions are mainly explained by the presence of repetitions in the set of unitigs. We find sufficient conditions for the existence of multiple solutions zones and propose an algorithm for identifying these zones.
- By using the optimal path found by the MILP model, our algorithm permits merging a set of unitigs satisfying the link distances into what we call *distance-based contigs*. These contigs, together with the other unitigs, are given to QUAST [6] for assessment.
- We tested this strategy on a set of 33 chloroplast genome data and compared the results with some of the most recent scaffolders (namely with SPAdes [1], SSPACE [2], BESST [15] and SWALO [14]).
- Our numerical experiments show that our approach produces assemblies of higher quality than the above heuristics on the considered benchmark.

## 2 Modeling the scaffolding problem

In this section we adapt the optimization approach proposed in [4] to the particularities and characteristics of the chloroplast genomes. Section 2.1 describes the graph modeling that is common for both approaches, while the mathematical programming formulation presented in section 2.2 includes enhancements of the model that, while making it less general, greatly increase its efficiency for chloroplast genome scaffolding.

### 2.1 Graph Modeling

The input data for our approach are the following:

- A set of *unitigs* together with their *repetition factor*. Unitigs represents unambiguous paths of a *de-Bruijn* graph. Only unitigs larger than a predefined threshold (cf section 4.1) are considered. The repetition factor is determined from k-mer counting techniques (cf section 4.1.2).
- A list of overlaps between the unitigs. Two unitigs overlap if they share a minimum of common nucleotides at their extremities.
- A list of oriented couples of unitigs (links). Links are determined from *paired-end* or *mate-pair* information. Due to insert size fluctuation, an interval distance is associated with any link from this list.

We follow the modeling from [4] where the scaffolding problem is reduced to a path finding in a directed graph *G* = (*V, E*), called a unitig graph, where both vertices *V* and edges *E* are weighted. The set of vertices *V* is generated based on the set *C* of the unitigs according the following rules: the unitig *i* is represented by at least two vertices *v_i_* and 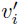 (forward/inverse orientation respectively). If the unitig i is repeated *k_i_* times (this value corresponds to the repetition factor), it generates a set *C_i_* of 2*k_i_* vertices. If two different vertices *v* and *w* belong to *C_i_* and have the same orientations, we can use the notation *v* ≈ *w*. Let us denote 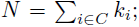 thus |*V*| = 2*N*.

The edges are generated following given patterns—a set of known overlaps/distances between the unitigs. Any edge is given in the graph *G* in its forward/inverse orientation. We denote by *e_ij_* the edge joining vertices *v_i_* and *V_j_* and the inverse of edge *e_ij_* by 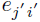. Let *w_v_* be the length of the unitig corresponding to vertex *v* and denote *W* = Σ_*v*∈*V^w_v_^*_. Moreover, let the weight *l_e_* on the edge *e* = (*v_i_,v_j_*) correspond to the value of the overlap/distance between unitigs represented by *v_i_* and *v_j_*. The problem then is to find a path in the graph *G* such that the total length (the sum over the traversed vertices and edges) is maximized, while a set of additional constraints are also satisfied:

- For any *i*, either vertex *v_i_* or 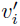 is visited (partici-pates in the path).
- The orientations of the nodes does not contradict the constraints imposed by the links. This is at least partially enforced by the construction of *G*.

To any edge *e* ∈ *E* we associate a variable *x_e_*. Its value is set to 1, if the corresponding edge participates in the assembled genome sequence (the associated path in our case), otherwise its value is set to 0. There are two kinds of edges: edges corresponding to overlaps between unitigs, denote them by *O* (from overlaps), and edges associated with the links relationships, denote them by *L*. We therefore have *E* = *L* ∪ *O*. Let *l_e_* be the length assigned to the edge *e* = (*u, v*). We define *l_e_* ∀*e* ∈ *O* such that *l_e_* < 0 and |*l_e_*| < min {*w_u_,w_v_*} is the overlap between the contigs corresponding to *v_i_* and *v_j_*, and 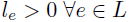, where *l_e_* is the link distance between unitigs represented by *v_i_* and *v_j_*.

Let 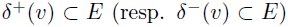 denote the sets of edges outgoing from (resp. incoming to) *v*.

### 2.2 Mixed Integer Linear Programming Formulation

The crucial observation in the approach proposed in [4] is that the genome can be assembled by searching for a particular longest path in the associated unitig graph. However, the beginning and the end of this path are unknown in the general case. This constraint leads to the sophisticated model described in [4]. Here we use the following two facts for chloroplast genomes in order simplify the above general approach:

1. Chloroplast genomes are circular;
2. One can safely assume that the largest unitig (say *s*) is always present in the solution.

Consequently, we introduce a supplementary vertex *t* that gets all incoming edges from *s*. Specifically, each edge (*x,s*) we replace by an edge (*x,t*) and set *δ*^−^(*t*) = *δ*^−^(*s*), *δ*^+^(*t*) = ∅, and *δ*^−^(*s*) = ∅. Vertices *s* and *t* will be considered respectively as the source (start) and the sink (end) of the path we are looking for.

Furthermore, to any vertex *v* ∊ *V* \ {*s*} we associate the variable *i_v_* s.t.

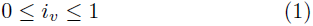

encoding whether *v* is in the solution path. Moreover, each vertex (or its inverse) should be visited at most once, which we encode as

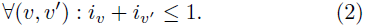

We associate a binary variable for any edge of the graph, i.e.,

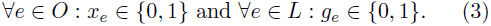

The two possibles states for a vertex *v* (to be (or not) an intermediate vertex in the path) are enforced by the following constraints

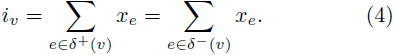

It is then obvious that the real variables *i_v_*, ∀*v* ∈ *V* take binary values.

We introduce a continuous variable *f_e_* ∈ *R*^+^ to express the quantity of the flow circulating along the edge *e* ∈ *E*. Without this variable, the solution found may contains some loops and hence may not be a simple path. We put a requirement that no flow can use an edge *e* when *x_e_* = 0, which can be encoded as

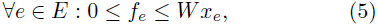

where *W* is as defined above (*W* = Σ_*v*∈*V*_ *w_v_*).

We use the flows *f_e_* in the following constraints, ∀*v* ∈ *V*\{*s*},

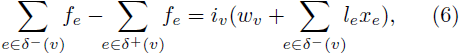

while for the source vertex we require

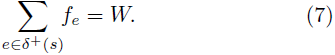

We furthermore observe that, because of (4), the constraint (6) can be written as follows

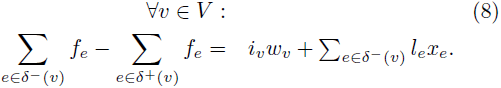

The constraint (8) is linear and we keep it in our model instead of (6).

The model so far defines a solution to the longest path problem. We need also to add information related to the links distances. For that reason, we associate a binary variable *g_e_* with each link *e*. For (*u,v*) ∈ *L*, the value of ∈*g*(*_u_,_v_*) is set to 1 only if both vertices *u* and *v* belong to the selected path and the length of the considered path between them is in the given interval 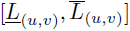. Constraints related to links are:

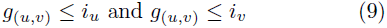

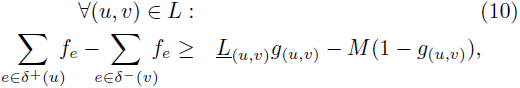

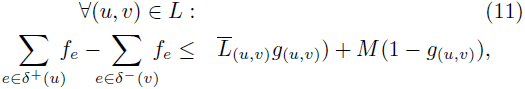

where *M* is some big constant.

Our goal is to find a long path in the graph such that as many as possible link distances are satisfied. The corresponding objective function hence is of the form

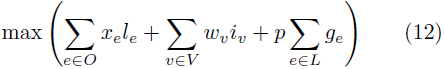

where *p* is a parameter to be chosen as appropriate (currently *p* = 1).

## 3 Dealing with multiple optimal solutions

By its nature, the information provided by the overlaps and mate pairs is not always sufficient to determine the assembly in a unique way. For instance, the unitig graph G is symmetric by constriction, e.g., if there is an edge (*v,W*) between vertices *v* and *W*, then there is an edge 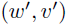 between their inverses 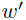 of *w* and 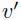 of *v*. Moreover, it contains repeated identical unitigs, which are modeled by different vertices of *G*. For all above reasons, for each optimal solution (path) *p^*^* found by our algorithm, there are typically multiple (exponential in the worst case) number of equivalent solutions (paths). Such paths are different from *p^*^* as sequences of vertices of *G*, but correspond to the same set of unitigs (and their inverted copies) and satisfy the same number of links, and hence are equally “optimal” from the point of view of the optimization problem (1)–(12). This issue is especially pronounced for chloroplasts due to their higher number of repeated/symmetrical regions.

Choosing just any arbitrary path from the set of equivalent optimal ones can result into an assembly different from the genome reference, which is the main criterion for evaluating the accuracy of the prediction. Therefore, our strategy is to detect in the optimal path multiple solutions portions and to separate them from subpaths that cannot be replaced be equivalent ones. This second type of subpaths will be merged in what we call *db-contigs* (contiguous sequences that satisfy the link distances). Obviously, none of the optimal solutions is eliminated while proceeding in such a manner. We call these zones “unsafe” and “safe,” respectively, and describe in this section a way to identify them.

Formally, we call two paths *p*_1_ = (*v_i_*,…, *v_k_*) and *P*_2_ = (*w*_1_,…, *W_k_*) of *G equivalent*, if they satisfy the same set of links and their components are permutations of the same set of unitigs (and their inverted copies). These paths can differ (or not) as sequences of base pairs. If *p* is a path in the unitig graph representing a solution of the optimization problem, we call a subpath 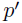 of *p* a *safe zone* of *p* if there exists no path in the graph *G* minus 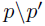 that is equivalent to and different from 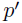 and 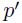 is a maximal subpath with this property. Safe zones are in fact subpaths containing a number of satisfied links, since each such link adds a constraint that reduces the number of subpaths that may be equivalent to it. Removing all safe zones from *p* leaves a set of paths that we call *unsafe zones*. We call a path *p link-closed* if for any link that has as an endpoint an intermediate vertex of *p*, its other endpoint is also *p*.

Next, we will illustrate a method for identifying unsafe zones by an example. Consider a unitig *v_s_* of multiplicity two. According to the graph-generation rules, there are vertices *v*_*s*0_ and *v*_*s*1_ in *G* corresponding to *v_s_* in the forward orientation and their corresponding vertices 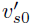 and 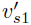 in the opposite direction. Assume also that there exists a link-closed subpath *p* = (*v_k_*, *v*_*k*+1_,…, *v_r_*) of a solution to the optimization problem such that *v_k_* = *v*_*s*0_ and 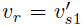. Remember that, for each edge (*v_i_*, *v*_*i*+1_) from *p*, the inverse edge 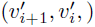 also exists in the unitig graph. Then we show that the inverse of *p*, i.e. the path 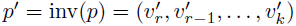 of inverted unitigs is also an optimal solution of the optimization problem. Obviously, 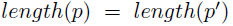. Since 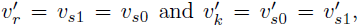 the paths *p* and 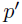 have the same sets of unitigs corresponding to their vertices and have identical unitigs at the beginning and their ends, but they are different as paths (sequences of vertices). The subsequence *p* = (*v*_*k*+1_,…, *v*_*r*−1_) is in this sense unsafe zone in the solution path. An example of such unsafe zone is illustrated on Figure 1.

**Figure 1:**
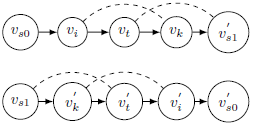
**Top**: a path *p* containing two links visualized with dashed lines; **Bottom**: its reversible path 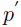. Note that *v*_*s*0_(resp. 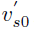) is identical to *v*_*s*1_(resp. 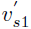).

It turns out that the type of subpath illustrated in the previous example is quite common and most of the unsafe zones that we have identified in our experiments can be captured using it. The algorithm for safe/unsafe zones detection based on using this pattern works as follows:

1. The vertices belonging to any satisfied link from the optimal path *p*^*^ found by the model in section
2. Potential db-contigs that overlap at least one vertex are merged in new (longer) potential db-contigs.
3. Any vertex outside the potential db-contigs is considered as unsafe.
4. For any potential db-contig *C* we apply the following algorithm.
  a. Any vertex *v_s_* ∈ *C* is initialized as safe.
  b. For any safe vertex *v_s_* ∈ *C* with multiplicity of at least two, and such that exists a couple 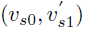 belonging to *C*, and such that the subpath between *v*_*s*0_ and 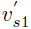 is link-closed do: (i) indicate as unsafe both vertices *v_s_*_0_ and 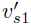; (ii) indicate the path between *v_s0_* and 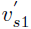 as a new potential db-contig.
5. All adjacent safe vertices are merged in true db-contigs (new meta-vertices).

The algorithm is illustrated on Figure 2, Figure 3 and Figure 4. In order to evaluate the quality of obtained solution we use QUAST [6]. Note that this tool requires for input just a set of contigs without indication for their repetition and orientation (for example, the input concerning the instance from Figure 4 consists in contigs *C*_1_, *C*_2_, *v*_1_ and *v*_80_ uniquely). QUAST maps any of them to the reference genome on order to assess its quality.

**Figure 2:**
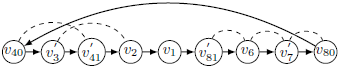
The initial solution.

**Figure 3:**
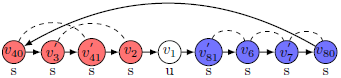
Steps 1, 2 and 3. Two potential db-contigs are created (the first one is red colored, the second is blue colored). Their vertices are initially labeled as safe. The vertex *v_1_* is labeled as unsafe since it is outside the potential db-contigs.

**Figure 4:**
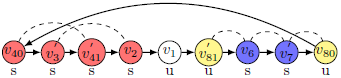
Step 4. Two repetitions are detected: the couples 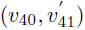 and 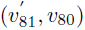. However, the path 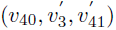 is not reversible, since it is not link-closed (because of the link 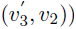. On the other hand, the path 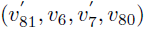 is reversible. The vertices 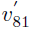 and *v*_80_ are labeled as unsafe. Finally, two true db-contigs are created: the first one, *C*_1_ (in red), contains the subpath 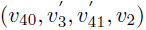, the second one *C_2_* (in blue), contains the subpath 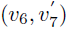. These two db-contigs, together with vertices/unitigs *v*_2_ (in white) and *v_80_* (in yellow) are given for assessment to QUAST.

Note that this algorithm does not necessarily find all unsafe/safe zones, but it works well in practice. Correctly identifying all such zones is an interesting research problem, whose solution can further improve the quality of our tool. In the next section we report some experimental results comparing our tool with some of the best existing similar tools.

The meta-vertices and the unsafe vertices, together with the edges that they induce, can be visualized as a meta-graph. Any cycle in this meta-graph is a potential genome assembly (see Figure 5 for illustration). However, only one of these cycles will be identical with the reference genome (if provided).

**Figure 5:**
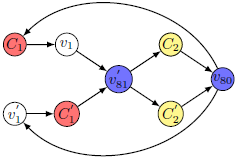
The metagraph obtained by the algorithm. Any cycle in this graph is an optimal solution and a potential genome assembly.

## 4 Experimental Analysis

### 4.1 Data Generation

#### 4.1.1 Simulated Data

From 33 chloroplast reference genomes (cf Table 4.1.4), 33 datasets of mate-pairs or pair-ended reads are generated with the art-illumina software with 100x depth of coverage [7]. For each dataset, the two following tasks are performed: (i) unitig generation; (ii) link computation.

#### 4.1.2 Unitig generation

Unitigs are generated with the Minia assembler [3]. A range of different *k*-mer sizes are tried to find the one that yields the best assembly.

For each unitig, its abundance (repetition factor) is computed, that is, the number of times it appears in the genome. For that, we define the kmer abundance as the number of times this kmer or its reverse-complement appears in the read files. The abundance of a unitig is then computed as the average abundance of all its kmers. This abundance is computed and returned by the Minia software.

In theory, the abundance of a unitig that is not repeated in the genome should be equal to the depth of coverage of sequencing, twice that amount for duplicated unitigs, and so on. We assume that the longest unitig is not duplicated, i.e. that its abundance is equal to the depth of coverage. The multiplicity of each unitig is then simply computed as its abundance divided by the depth of coverage, rounded to the nearest upper integer value.

This strategy provides an estimation of the coverage, but its accuracy strongly depends of the length of the unitigs. Longer the unitigs, better the estimation. Actually, for very short unitigs, we can only provide intervals of confidence or, at least an upper bound.

#### 4.1.3 Link computation

Each mate-pair or pair-ended read is individually mapped to unitigs with minimap [10]. We discard reads that map ambiguously to several locations. Reads of a pair that map to different unitigs indicate a mate-pair link in the graph. To avoid false positives, we only keep links that are validated by at least 5 pairs. The link size is estimated thanks to the known inserts size and mapping position in each unitig, and averaged over all pairs that confirm the link.

#### 4.1.4 Computational results

We have generated a data set of 33 chloroplasts genomes obtained from the NCBI website (https://www.ncbi.nlm.nih.gov/genome). In order to simulate mate-pairs and pair-ends sequencing, we used the ART simulator Illumina [8]. We have produced reads with a length of 250bp and 100X coverage. Two types of simulation were performed: for the pair-end simulation the inserts size was 600bp, while, for the mate-pairs simulation, we used an insert of 8000bp. The reads were subsequently assembled in unitigs by Minia [3] and the generated *fasta* file was input to the scaffolders that needed it (SPAdes [1] and SWALO [14] work directly with the reads and do not require it). The unitigs were produced with an abundance of 4 and a *k*-mer of 125.

The assemblies were evaluated by QUAST [6] tool by comparison with the reference genome that was used for the simulation. (More detailed experimental data is given in the Appendix.) Our tool is denoted by GAT (Genscale Assembly Tool). In our experiments, GAT has been as good as, and often better, than the best current scaffolding tools, while ensuring good coverage of the reference genome (a parameter that tends to degrade with other scaffolders). It has been particularly good in case of pair-ends computations by ensuring a regular and nearly optimal assembly.

During the mate pairs computations GAT performed well by producing often the smallest number of contigs and was outperformed only in the case of Atropa genome. Figure 6 illustrates that GAT produces on average fewer contigs than its competitors. Moreover, it ensures the best genome coverage as we can observe on Figure 7. This indicates that the output produced by our tool are reliable, complete, and don’t lose information compared to the original genome. SWALO failed to assemble 10 genomes out of 33, SSPACE 3 genomes, and BESST-one genome. SPADES and GAT where the only tools for which QUAST did not indicate any missassemblies.

**Figure 6:**
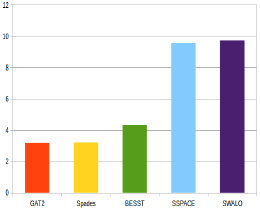
Mate-pairs data: Average number of contigs comparison.

**Figure 7:**
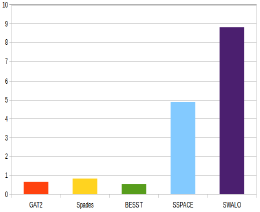
Mate-pairs data: Average fraction of genome left out comparison.

In the case of pair-end simulations, we have obtained equally good results. The performance of GAT and SPAdes are very close in term of average number of contigs (cf. Figure 8). However, SPAdes is clearly outperformed by GAT, BESST and SWALO concerning the genome coverage (cf. Figure 9). On this figure we also observe that BESST is as reliable as GAT, but it couldn’t solve Euglena (21)-something that GAT achieved.

**Figure 8:**
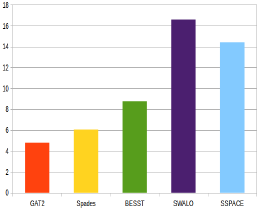
Pair-ends data: Average number of contigs comparison.

**Figure 9:**
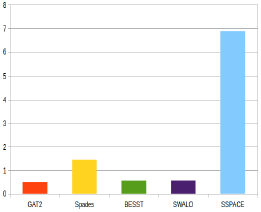
Pair-ends data: Average fraction of genome left out comparison.

**Figure 10:**
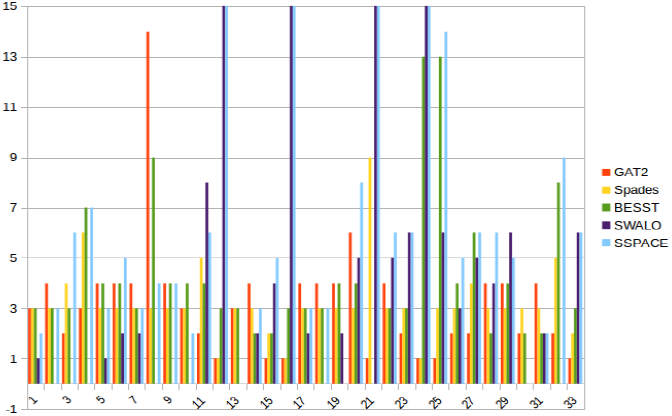
Mate-pairs data: number of contigs comparison between Spades, Sspace, Besst, Swalo and GAT.

**Figure 11:**
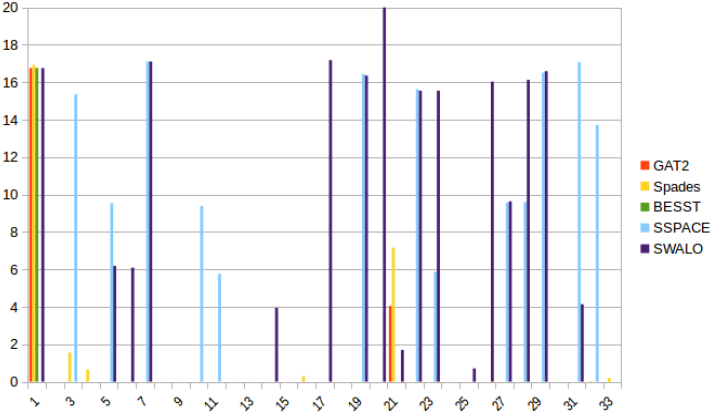
Mate-pairs data: fraction of genome left out comparison between Spades, Sspace, Besst,Swalo and GAT.

**Figure 12:**
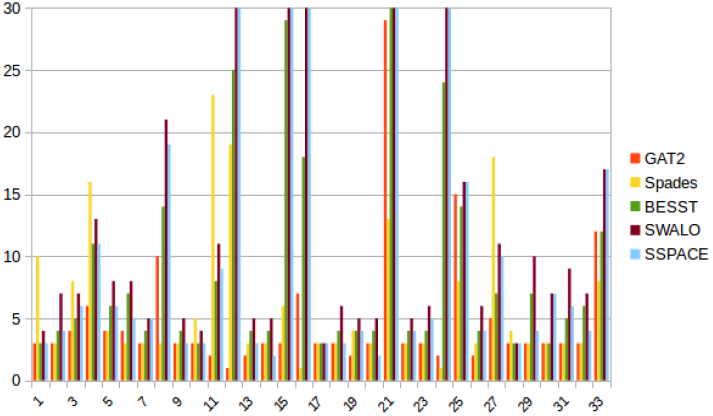
Pair-ends data: number of contigs comparison between Spades, Sspace, Besst, SWALO and GAT.

**Figure 13:**
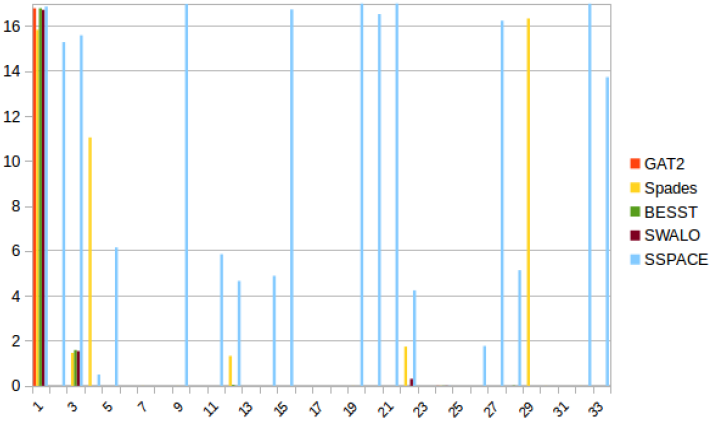
Pair-ends data: fraction of genome left out comparison between Spades, Sspace, Besst, SWALO and GAT.

**Table 1.** The benchmark containing 36 chloroplast genomes whose names given in the first column. The second column contains their lengths. We observed that this value equals the value given by the first term of the objective function (12). The third and fourth columns give the size of the graph (i.e. number of vertices and edges). |*L*| indicates the number of given links, while NSL stands for number of satisfied links in the solution.

| No | Genomes | Size | \|V\| | \|O\| | \|L\| | NSL |
| --- | --- | --- | --- | --- | --- | --- |
| 1 | Acorus Calamus | 153821 | 8 | 16 | 16 | 3 |
| 2 | AdiantumCapillus Veneris | 150568 | 20 | 24 | 24 | 5 |
| 3 | Agrostis Stolonifera | 136584 | 20 | 52 | 24 | 6 |
| 4 | Angiopteris Evecta | 153901 | 34 | 78 | 70 | 12 |
| 5 | Anthoceros Formosae | 161162 | 16 | 32 | 24 | 5 |
| 6 | Arabidopsis Thaliana | 161162 | 20 | 40 | 32 | 7 |
| 7 | Arabishirsuta | 153689 | 12 | 24 | 24 | 5 |
| 8 | Atropa | 156687 | 46 | 90 | 34 | 9 |
| 9 | Capsella Bursa Pastoris | 154490 | 12 | 24 | 24 | 5 |
| 10 | Chaetosphaeridium Globosum | 131183 | 8 | 16 | 16 | 3 |
| 11 | Chara Vulgaris | 184933 | 24 | 56 | 24 | 7 |
| 12 | Chlorella Vulgaris | 150613 | 52 | 50 | 50 | 24 |
| 13 | Chlorokybus Atmophyticus | 152229 | 10 | 18 | 18 | 4 |
| 14 | Citrus Sinensis | 160129 | 12 | 24 | 24 | 5 |
| 15 | Cyanidioschyzon Merolae | 149067 | 72 | 82 | 46 | 22 |
| 16 | Cyanidium Caldarium | 164921 | 38 | 36 | 32 | 15 |
| 17 | Daucus Carota | 155911 | 8 | 16 | 16 | 3 |
| 18 | Draba Nemorosa | 153289 | 12 | 24 | 24 | 5 |
| 19 | Eimeria Tenella | 160604 | 10 | 18 | 18 | 4 |
| 20 | Epifagus Virginiana | 70028 | 12 | 24 | 24 | 5 |
| 21 | Euglena Gracilis | 143171 | 146 | 554 | 30 | 5 |
| 22 | Gossypium Barbadense | 160317 | 12 | 24 | 24 | 5 |
| 23 | Gossypium Hirsutum | 160301 | 14 | 28 | 24 | 5 |
| 24 | Gracilaria Tenuistipitata | 183883 | 54 | 54 | 44 | 21 |
| 25 | Guillardia Theta | 121524 | 44 | 88 | 24 | 5 |
| 26 | Helianthus Annuus | 151104 | 10 | 18 | 18 | 4 |
| 27 | Huperzia Lucidula | 154259 | 20 | 48 | 20 | 5 |
| 28 | Lactuca Sativa | 152765 | 8 | 16 | 16 | 3 |
| 29 | Lepidium Virginicum | 154743 | 24 | 48 | 48 | 11 |
| 30 | Liriodendron Tulipifera | 159886 | 8 | 16 | 16 | 3 |
| 31 | Lobularia Maritima | 152659 | 16 | 32 | 32 | 7 |
| 32 | Lotus Corniculatus | 150519 | 20 | 80 | 32 | 7 |
| 33 | Pinus | 116864 | 58 | 128 | 12 | 6 |

## 5 Conclusion

Here we design and test an algorithm for scaffolding and gap filling phases in the case of circular genomes. Our approach is based on a version of the longest path problem solved by MILP modeling. It works both in case of mate-pairs and pair-ends distances. On a benchmark of 33 chloroplast genomes our algorithm significantly outperforms four recent scaffolding heuristics with respect to the quality of the scaffolds. The obtained results fully justify the efforts for designing exact approaches for genome assembly. Regardless of that, we consider the current results as a work in progress. The biggest challenge is to extend the method to much bigger genomes. We are currently implementing advanced combinatorial optimization decomposition techniques to increase the scalability of the approach without sacrificing the accuracy of the results.

## Acknowledgments

We would like to thank Guillaume Rizk for adapting the Minia assembler [3] for the purpose of our algorithm and for his help in data generation. Many thanks to Rayan Chikhi for valuable discussions.

## References

[1] Anton Bankevich, Sergey Nurk, Dmitry Antipov, Alexey A Gurevich, Mikhail Dvorkin, Alexander S Kulikov, Valery M Lesin, Sergey I Nikolenko, Son Pham, Andrey D Prjibelski, Alexey V Pyshkin, Alexander V Sirotkin, Nikolay Vyahhi, Glenn Tesler, Max A Alekseyev, and Pavel A Pevzner. Spades: a new genome assembly algorithm and its applications to single-cell sequencing. Journal of computational biology: a journal of computational molecular cell biology, 19(5):455–477, May 2012.

[2] Marten Boetzer, Christiaan V. Henkel, Hans J. Jansen, Derek Butler, and Walter Pirovano. Scaffolding pre-assembled contigs using SSPACE. Bioinformatics (Oxford, England), 27(4):578–579, February 2011.

[3] Rayan Chikhi and Guillaume Rizk. Space-efficient and exact de bruijn graph representation based on a bloom filter. Algorithms for Molecular Biology, 8(1):22, 2013.

[4] Sebastien François, Rumen Andonov, Hristo Djidjev, and Dominique Lavenier. Global optimization methods for genome scaffolding. In 8th International Network Optimization Conference (INOC), 2017. to appear in the special issue of Electronic Notes in Discrete Mathematics (ENDM) V. 64.

[5] Alexey A. Gritsenko, Jurgen F. Nijkamp, Marcel J.T. Reinders, and Ridder Dick de. GRASS: a generic algorithm for scaffolding next-generation sequencing assemblies. Bioinformatics, 28(11):1429–1437, 2012.

[6] Alexey Gurevich, Vladislav Saveliev, Nikolay Vyahhi, and Glenn Tesler. Quast: quality assessment tool for genome assemblies. Bioinformatics, 29(8):1072–1075, 2013.

[7] Weichun Huang, Leping Li, Jason R Myers, and Gabor T Marth. Art: a next-generation sequencing read simulator. Bioinformatics, 28(4):593–594, 2011.

[8] Weichun Huang, Leping Li, Jason R. Myers, and Gabor T. Marth. Art: a next-generation sequencing read simulator. Bioinformatics, 28(4):593–594, 2012.

[9] Daniel H. Huson, Knut Reinert, and Eugene W. Myers. The greedy path-merging algorithm for contig scaffolding. J. ACM, 49(5):603–615, 2002.

[10] Heng Li. Minimap and miniasm: fast mapping and de novo assembly for noisy long sequences. Bioinformatics, 32(14):2103–2110, 2016.

[11] Paul Medvedev, Son Pham, Mark Chaisson, Glenn Tesler, and Pavel Pevzner. Paired de Bruijn graphs: A novel approach for incorporating mate pair information into genome assemblers. Journal of Computational Biology, 18(11):1625–1634, 11 2011.

[12] Briot Nicolas, Chateau Annie, Rémi Coletta, Simon de Givry, Philippe Leleux, and Schiex Thomas. An integer linear programming approach for genome scaffolding. In Workshop on Constraint based Methods for Bioinformatics, 2015.

[13] Pavel A. Pevzner, Haixu Tang, and Michael S. Waterman. An Eulerian path approach to DNA fragment assembly. PNAS, 98(17):9748–9753, 2001.

[14] Atif Rahman and Lior Pachter. Swalo: scaffolding with assembly likelihood optimization. bioRxiv, 2016.

[15] Kristoffer Sahlin, Francesco Vezzi, Björn Nystedt, Joakim Lundeberg, and Lars Arvestad. BESST - efficient scaffolding of large fragmented assemblies. BMC Bioinformatics, 15:281, 2014.

[16] James L. Weber and Eugene W. Myers. Human whole-genome shotgun sequencing. Genome Research, 7(5):401–409, 1997.

[17] Mathias Weller, Annie Chateau, and Rodolphe Giroudeau. Exact approaches for scaffolding. BMC bioinformatics, 16(Suppl 14):S2, 2015.

